# NCOA4 drives ferritin phase separation to facilitate macroferritinophagy and endosomal microferritinophagy

**DOI:** 10.1101/2022.01.31.478434

**Authors:** Tomoko Ohshima, Hayashi Yamamoto, Yuriko Sakamaki, Chieko Saito, Noboru Mizushima

**Affiliations:** Department of Biochemistry and Molecular Biology, Graduate School and Faculty of Medicine, The University of Tokyo, Tokyo 113-0033, Japan; Microscopy Research Support Unit, Research Core, Tokyo Medical and Dental University, Tokyo 113-8519, Japan

## Abstract

A ferritin particle consists of 24 ferritin proteins (FTH1 and FTL) and stores iron ions within it. During iron deficiency, ferritin particles are transported to lysosomes to release iron ions. Two transport pathways have been reported: macroautophagy and ESCRT-dependent endosomal microautophagy. Although the membrane dynamics of these pathways differ, both require NCOA4, which is thought to be an autophagy receptor for ferritin. However, the exact function of NCOA4 remains elusive. Here, we found that ferritin particles form liquid-like condensates in a NCOA4-dependent manner. Homodimerization of NCOA4 and interaction between FTH1 and NCOA4 (i.e., multivalent interactions between ferritin particles and NCOA4) were required for the formation of ferritin condensates. Disruption of these interactions impaired ferritin degradation. Time-lapse imaging and three-dimensional correlative light and electron microscopy revealed that these ferritin–NCOA4 condensates were directly engulfed by autophagosomes and endosomes. In contrast, TAX1BP1 was not required for the formation of ferritin–NCOA4 condensates but was required for their incorporation into autophagosomes and endosomes. These results suggest that NCOA4 acts not only as a canonical autophagy receptor but also as a driver to form ferritin condensates to facilitate the degradation of these condensates by macroautophagy (i.e., macroferritinophagy) and endosomal microautophagy (i.e., microferritinophagy).

## Introduction

Iron is essential for various biological processes, but in excess it can cause deleterious effects (Pantopoulos et al., 2012; Bogdan et al., 2016; Fenton, 1894; Haber and Weiss, 1934; Halliwell and Gutteridge, 1990). Thus, the intracellular iron level must be tightly regulated. Ferritin is a key player in the regulation of iron homeostasis and is highly conserved among organisms except for some fungi (e.g., yeasts) and several bacterial and archaeal species (Canessa and Larrondo, 2013; Bai et al., 2015). It forms a particle consisting of 24 subunits of ferritin heavy chain 1 (FTH1) and ferritin light chain (FTL), which incorporates up to 4,500 Fe^2+^ ions, oxidizes them to Fe^3+^, and stores them as mineral cores in its hollow cavity (Mann et al., 1986). During iron deficiency, ferritin particles are transported to lysosomes in order to release the stored iron atoms (Kidane et al., 2006; Asano et al., 2011). Ferritin delivery to lysosomes is mediated by two different pathways: macroautophagy (Asano et al., 2011; Mancias et al., 2014; Dowdle et al., 2014; Goodwin et al., 2017) and the endosomal sorting complexes required for the transport (ESCRT)-mediated pathway (Goodwin et al., 2017). To degrade ferritin, both pathways require nuclear receptor coactivator 4 (NCOA4), currently considered to be a ferritin receptor, and the macroautophagy adaptor Tax1-binding protein 1 (TAX1BP1) (Mancias et al., 2014; Dowdle et al., 2014; Goodwin et al., 2017). Because NCOA4 and TAX1BP1 are degraded by not only macroautophagy but also endosomal microautophagy (Mejlvang et al., 2018), the ESCRT-dependent ferritin degradation reported by Goodwin *et al*. (Goodwin et al., 2017) is likely mediated by microautophagy. Although NCOA4 has been shown to interact with TAX1BP1 (Goodwin et al., 2017), little is known about how it is involved in ferritin degradation through the morphologically and mechanistically distinct pathways of macroautophagy and endosomal microautophagy.

Electron microscopy studies in cultured cells and tissues have demonstrated that ferritin particles can be clustered in the cytosol (Heynen and Verwilghen, 1982; Takano-Ohmuro et al., 2000) as well as within membranous structures (Sullivan et al., 1976; Heynen and Verwilghen, 1982; Iancu et al., 2014). Furthermore, our group previously found that clusters of ferritin particles are enlarged in autophagy-deficient cells, some of which are over 500 nm in size, and accumulate at autophagosome formation sites (Kishi-Itakura et al., 2014). These ferritin clusters are spherical and often associate with SQSTM1 (also called p62) bodies without mixing their contents. Based on these findings, we hypothesized that ferritin clusters are liquid-like condensates caused by liquid–liquid phase separation (LLPS).

In this study, we show that ferritin clusters indeed have liquid-like properties and that NCOA4 is required for ferritin phase separation. The ferritin–NCOA4 condensates are incorporated into autophagosomes and endosomes in a TAX1BP1-dependent manner. Failure of condensate formation impairs ferritin degradation. These data suggest that the formation of liquid-like ferritin–NCOA4 condensates is a common mechanism to facilitate degradation by both macroautophagy and endosomal microautophagy.

## Results

### Ferritin particles assemble to form large condensates that exhibit liquid-like properties

To observe the distribution of ferritin particles in living cells, we established HeLa cells stably expressing monomeric EGFP (mGFP)-tagged FTH1 and FTL. Both mGFP-FTH1 and mGFP-FTL formed punctate structures in the cytoplasm in wild-type (WT) cells under normal (Figure 1A) and iron-replete conditions that induced ferritin expression (Figure 1B). The expression levels of mGFP-FTH1 and mGFP-FTL were comparable to (or even lower than) those of endogenous FTH1 and FTL (Figure 1C), suggesting that the formation of puncta was not due to overexpression. The ferritin puncta became larger in autophagy-deficient *FIP200* KO cells (Figures 1A, 1B). Transmission electron microscopy (TEM) of *FIP200* KO cells expressing GFP-FTH1 showed cytosolic spherical clusters of the electron-dense ferritin particles (Figure 1D), consistent with our previous report (Kishi-Itakura et al., 2014). In addition, the clustered ferritin structures were not enclosed by membranes (Figure 1D, right panel). These structural characteristics (i.e., the highly spherical shape and the lack of membranes) raised the possibility that the ferritin clusters are biomolecular condensates driven by LLPS. Fluorescence recovery after photobleaching (FRAP) measurements in WT cells revealed that mGFP-FTH1 condensates exhibited approximately 20% recovery within 10 min (Figures 1E, 1F). In addition, coalescence of two discrete GFP-FTH1 condensates was observed (Figure 1G). These features, together with the abovementioned structural characteristics, are consistent with the current criteria for liquid-like biomolecular condensates formed via LLPS (Alberti et al., 2019). Thus, these results suggest that ferritin particles have the propensity to congregate to form liquid-like biomolecular condensates.

**Figure 1.**
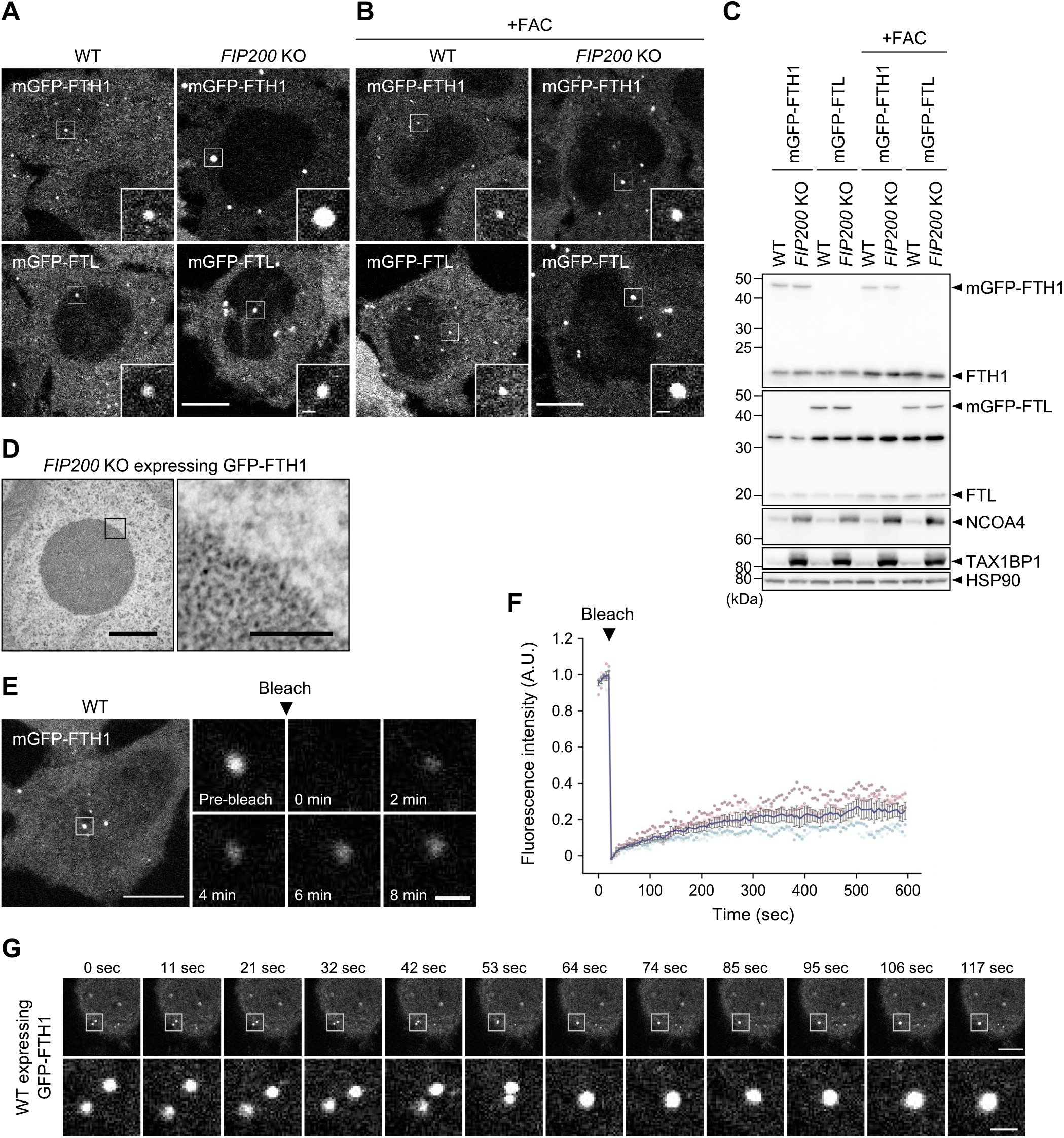
Ferritin particles assembled to form liquid-like condensates. **(A, B)** Fluorescent images of the ferritin subunits FTH1 and FTL. Wild-type (WT) and autophagy-deficient *FIP200* KO HeLa cells stably expressing mGFP-FTH1 or mGFP-FTL were grown in DMEM (A) or treated with 10 μg/mL ferric ammonium citrate (FAC) for 24 h **(B)** and observed by fluorescence microscopy. Scale bars, 10 μm (main) and 1 μm (inset). **(C)** Immunoblots showing the expression levels of mGFP-FTH1 and mGFP-FTL in cells used in (A). **(D)** TEM images of the ferritin condensates in *FIP200* KO HeLa cells expressing GFP-FTH1. Scale bars, 500 nm (main) and 100 nm (magnified). **(E)** FRAP analyses of the ferritin condensates. WT HeLa cells expressing mGFP-FTH1 were treated with 100 μg/mL FAC for 24 h followed by 50 μM deferoxamine (DFO) for 5 h, and subjected to FRAP analyses. Scale bars, 10 μm (main) and 1 μm (magnified). **(F)** Quantification of (E). Fluorescence intensities before photobleaching were set to 1. Representative results from two independent experiments are presented as means ± SEM (*n* = 5). **(G)** Coalescence of the ferritin condensates. WT HeLa cells expressing GFP-FTH1 were grown in DMEM and observed by time-lapse fluorescence microscopy at ∼11-s intervals. Scale bars, 10 μm (main) and 1.5 μm (magnified).

### NCOA4 drives ferritin phase separation

NCOA4 is involved in ferritin delivery to lysosomes by direct interaction with FTH1 (Mancias et al., 2014; Dowdle et al., 2014; Mancias et al., 2015). Thus, we investigated the relationship between ferritin condensates and NCOA4. In WT and *FIP200* KO cells, mGFP-NCOA4 co-localized with the mRuby3-FTH1 puncta (Figure 2A), indicating that NCOA4 is a component of ferritin condensates. Then, to investigate the role of NCOA4 in condensate formation, we generated *NCOA4* KO cells by using the CRISPR-Cas9 method (Figure S1A). The resulting cells showed a defect in the degradation of FTH1 upon iron depletion, achieved via the iron chelator deferoxamine (DFO) (Figure S1B), which was in line with previous reports (Mancias et al., 2014; Dowdle et al., 2014). Fluorescence microscopy revealed that mGFP-FTH1, which formed punctate structures in WT and *FIP200* KO cells, became diffuse in the absence of NCOA4 (Figure 2B), suggesting that NCOA4 is indispensable for the formation of ferritin condensates.

**Figure 2.**
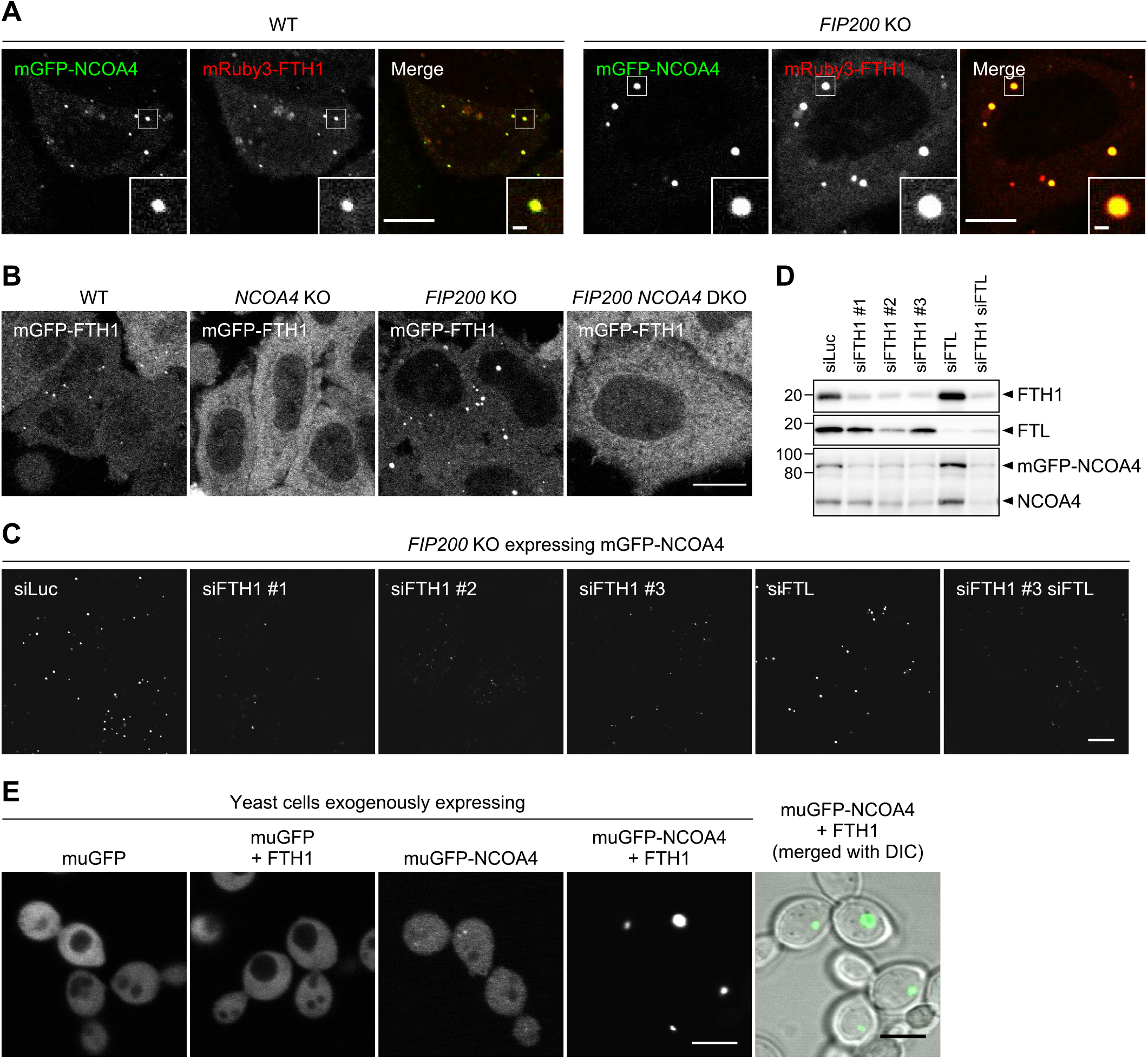
NCOA4 drives ferritin condensate formation. **(A)** Localization of NCOA4 in ferritin condensates. WT and *FIP200* KO cells expressing both mGFP-NCOA4 and mRuby3-FTH1 were grown in DMEM and observed by fluorescence microscopy. Scale bars, 10 μm (main) and 1 μm (inset). **(B)** NCOA4 is essential for condensate formation. WT, *NCOA4* KO, *FIP200* KO, and *FIP200 NCOA4* double KO (DKO) cells expressing mGFP-FTH1 were observed as in (A). Scale bar, 15 μm. **(C)** Knockdown of ferritin subunits. *FIP200* KO cells expressing mGFP-NCOA4 were transfected with the indicated siRNAs (siLuc, siFTH1, and siFTL) and observed by fluorescence microscopy. Scale bar, 10 μm. **(D)** Immunoblots showing the expression levels of FTH1, FTL, and NCOA4 in cells used in (C). **(E)** NCOA4 and FTH1 are sufficient for condensate formation. Yeast cells exogenously expressing muGFP-NCOA4 with or without FTH1 were observed by fluorescence microscopy. Scale bar, 5 μm.

Next, we examined whether FTH1 and FTL were necessary for the formation of ferritin–NCOA4 condensates. Knockdown of FTH1 but not FTL impeded mGFP-NCOA4 puncta formation in *FIP200* KO cells (Figures 2C, 2D). Although this result suggests that FTH1, but not FTL, is required for the formation of NCOA4 condensate, knockdown of FTH1 also reduced the expression levels of mGFP-NCOA4, which made it difficult to evaluate the specific role of FTH1 (Figures 2C, 2D).

Given that FTH1 interacts with NCOA4 (Mancias et al., 2015), we further investigated whether FTH1 and NCOA4 were sufficient for the formation of ferritin condensates. When muGFP-tagged human NCOA4 was exogenously expressed in yeast cells (ferritin and NCOA4 homologs are not present in yeast), it mostly dispersed in the cytoplasm and occasionally formed some small dots. In contrast, when co-expressed with FTH1, muGFP-NCOA4 formed one large punctum per cell (Figure 2E). These results suggest that FTH1 and NCOA4 are sufficient for the formation of ferritin–NCOA4 condensates.

### Ferritin–NCOA4 condensate formation is driven by NCOA4 self-interaction and NCOA4-FTH1 interaction

NCOA4 consists of the N-terminal coiled-coil domain (N), middle domain (M), and C-terminal domain (C), which are interconnected with intrinsically disordered regions (IDRs) (IDR1 and IDR2) (Figure 3A). To determine which domain(s) of NCOA4 is required for the formation of ferritin condensates, we constructed domain truncation mutants (Figure 3A). *NCOA4* KO cells expressing 3×FLAG-tagged NCOA4 (FLAG-NCOA4) restored the formation of mGFP-FTH1 puncta (Figures 3B, 3C). The ΔM and ΔC mutants of FLAG-NCOA4 also recovered the formation of puncta, and the ΔIDR1 mutant partially recovered it. In contrast, the ΔN and ΔIDR2 mutants failed to form mGFP-FTH1 puncta (Figures 3B, 3C). These results indicate the importance of the N and IDR2 domains of NCOA4 in the formation of ferritin condensates, which could be partly explained by the interaction of IDR2 with FTH1 (Mancias et al., 2015; Gryzik et al., 2017).

**Figure 3.**
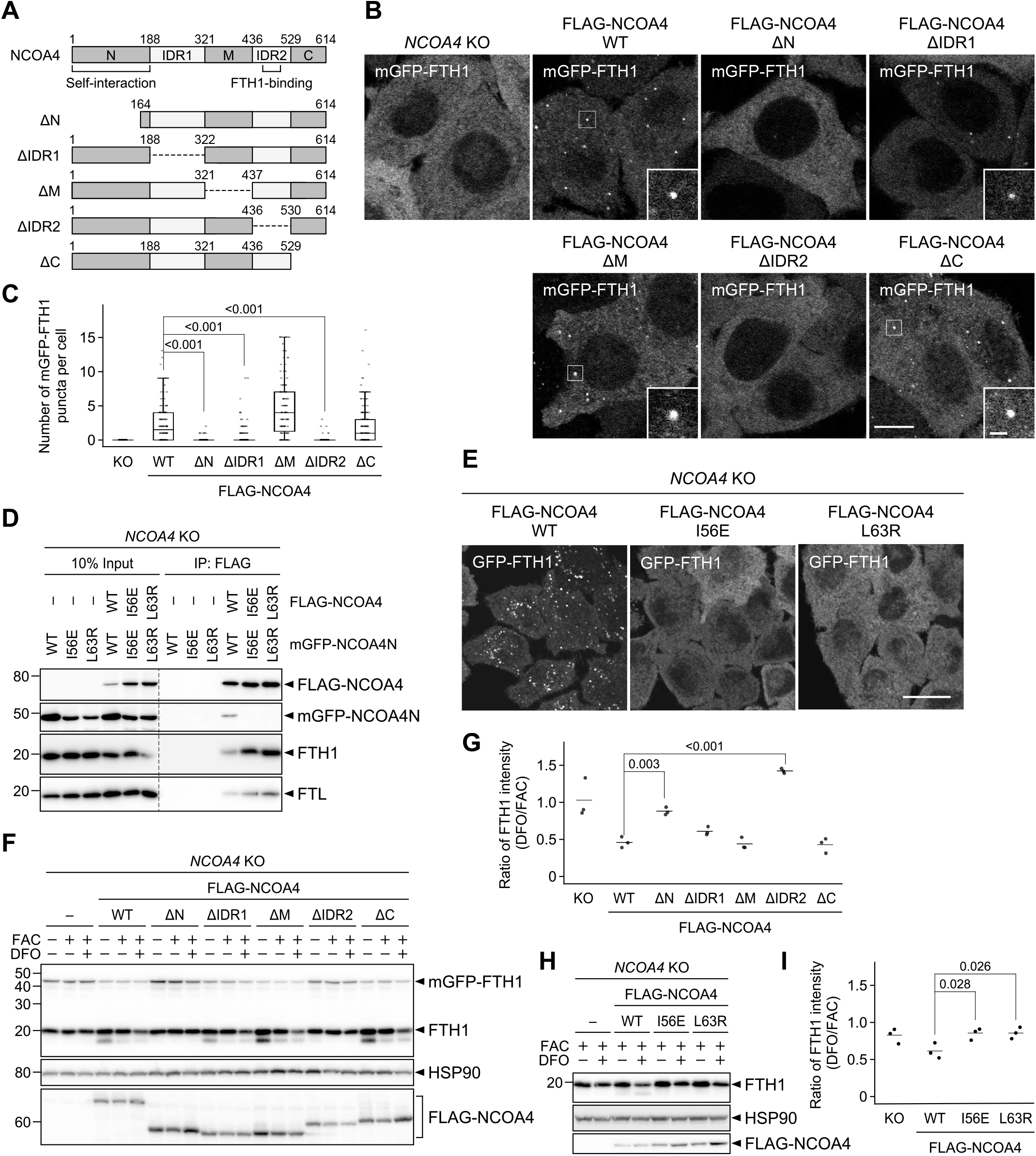
Ferritin–NCOA4 condensate formation is driven by NCOA4 self-interaction and responsible for ferritin degradation. **(A)** Truncated constructs of NCOA4. IDR, intrinsically disordered region. **(B)** WT or the truncated mutants of FLAG-NCOA4 were stably expressed in *NCOA4* KO cells harboring mGFP-FTH1. Scale bars, 10 μm (main) and 1.5 μm (inset). **(C)** Numbers of mGFP-FTH1 puncta per cell in (B) were counted (*n* = 269–344 cells from three biological replicates). Solid bars indicate the medians, boxes the interquartile ranges, and whiskers the 10th to 90th percentiles. Differences among the cells expressing FLAG-NCOA4 were statistically analyzed by Dunnett multiple comparison test. **(D)** Defective self-interaction of the N-terminal domain mutants of NCOA4. *NCOA4* KO cells expressing both FLAG-NCOA4 (full-length) and mGFP-NCOA4N (the N-terminal domain of NCOA4) with or without I56E or L63R mutation were subjected to co-immunoprecipitation using anti-FLAG antibody. The samples were analyzed by immunoblotting with the antibodies against FLAG, GFP, FTH1, and FTL. **(E)** The I56E or L63R mutant of FLAG-NCOA4 were stably expressed in *NCOA4* KO cells harboring GFP-FTH1. The cells were observed as in (A). **(F)** Cells used in (B) were grown in DMEM and treated with 10 μg/mL FAC for 24 h followed by 50 μM DFO for 12 h. Whole-cell lysates were analyzed by immunoblotting with the antibodies against FTH1, HSP90, and FLAG. **(G)** FTH1 degradation upon DFO treatment in (F). The ratio of the FTH1 band intensities under DFO-treated conditions to those under FAC-treated conditions is shown. Solid bars indicate the medians, and dots the data from three independent experiments. Differences were statistically analyzed by Dunnett multiple comparison test. **(H)** The cells used in (E) were examined as in (F). **(I)** Quantification of (H) as in (G).

The N-terminal coiled-coil domain of NCOA4 is known as a self-oligomerization domain (Monaco et al., 2001). In fact, full-length FLAG-NCOA4 interacted with the N-terminal domain of NCOA4 (NCOA4N) fused with mGFP (Figure 3D). The HHpred search (Soding et al., 2005) identified that the N-terminal coiled-coil domain in NCOA4 is structurally similar to that in TRIM28. TRIM28 forms a homodimer (Stoll et al., 2019), and Ile-299 and Leu-306 are located at the homodimerization interface (Figure S2). Thus, we introduced mutations in the corresponding residues in NCOA4 (I56E and L63R), which were predicted via trRosetta (Yang et al., 2020) (Figure S2). These mutations blocked self-interaction via the N-terminal domain (Figure 3D). The yeast two-hybrid assay using NCOA4N also showed that the I56E or L63R mutants were defective in self-interaction (Figure S3). Fluorescence microscopy revealed that the I56E or L63R mutants did not show GFP-FTH1 puncta in *NCOA4* KO cells (Figure 3E), indicating that NCOA4 self-interaction is required for the formation of ferritin–NCOA4 condensates. Taken together, we concluded that the formation of ferritin–NCOA4 condensates is driven by NCOA4-mediated multivalent interactions (i.e., NCOA4-FTH1 interaction and NCOA4 self-interaction).

### Ferritin–NCOA4 condensate formation is required for ferritin degradation

We then determined whether condensate formation was required for the degradation of ferritin. When introduced into *NCOA4* KO cells, WT and all the NCOA4 mutants that restored the formation of mGFP-FTH1 puncta (ΔM, ΔC, and ΔIDR1) also rescued FTH1 degradation upon iron depletion, whereas the ΔN and ΔIDR2 mutants, which failed to form mGFP-FTH1 puncta, also failed to degrade FTH1 (Figures 3F, 3G). Furthermore, the dimerization-defective NCOA4 mutants (I56E and L63R) failed to restore FTH1 degradation (Figures 3H, 3I). It should be noted that these I56E and L63R mutants retained the binding ability to FTH1 and FTL (likely through FTH1 in ferritin particles) (Figure 3D). Thus, these results suggest that condensate formation rather than NCOA4–ferritin binding itself is important for ferritin degradation.

### Ferritin–NCOA4 condensates are common substrates for macroautophagy and endosomal microautophagy

Next, we investigated whether ferritin–NCOA4 condensates were targeted by autophagosomes. WT cells expressing mGFP-FTH1 and HaloTag7-tagged LC3B (Halo-LC3), an autophagosomal membrane marker (Kabeya et al., 2000; Mizushima, 2004), were observed under iron-deficient conditions. Time-lapse fluorescence microscopy showed that some mGFP-FTH1 puncta were sequestered by cup-shaped autophagosomal membranes in a piecemeal manner (Figure 4A and Movie S1). To obtain more detailed morphological information, we conducted three-dimensional correlative light and electron microscopy (3D-CLEM). In line with the observations under a fluorescence microscope, we observed an electron-dense ferritin condensate with a diameter of ∼500 nm being sequestered partly by a Halo-LC3-positive autophagosomal membrane (Figure 4B). The autophagosomal membrane appeared to be in close contact with the ferritin condensate, similar to the fluidophagy of the SQSTM1 bodies (Agudo-Canalejo et al., 2021). The remaining region not engulfed by the autophagosomal membrane retained a spherical shape, consistent with its liquid-like property. These results suggest that ferritin–NCOA4 condensates are selective substrates for macroautophagy.

**Figure 4.**
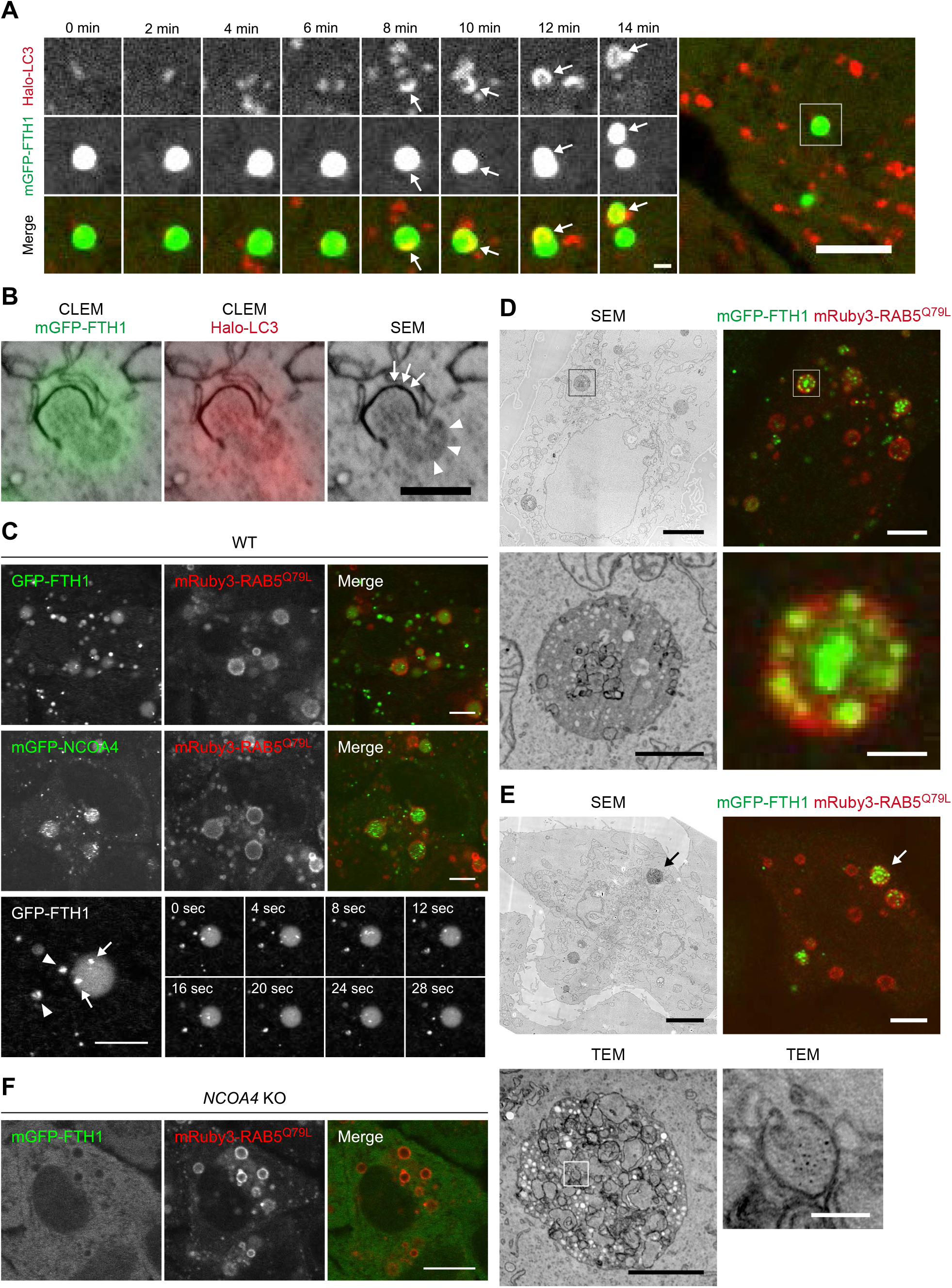
The ferritin–NCOA4 condensates are targeted by both macroautophagy and endosomal microautophagy. **(A)** Ferritin–NCOA4 condensates are engulfed by autophagosomes. WT cells expressing mGFP-FTH1 (green) and Halo-LC3 (red) were treated with 50 μg/mL FAC for 24 h followed by 50 μM DFO for 5 h and then observed by time-lapse fluorescence microscopy at 2-min intervals (see also Movie S1). Arrows indicate part of ferritin condensates engulfed by an autophagosome. Scale bars, 5 μm (main) and 1 μm (magnified). **(B)** 3D-CLEM of cells treated as in (A). Arrowheads indicate the surface of a ferritin–NCOA4 condensate exposed to the cytosol, and arrows an elongating autophagosomal membrane. Scale bar, 500 nm. **(C)** The ferritin–NCOA4 condensates are incorporated into endosomes. WT cells expressing GFP-FTH1 or mGFP-NCOA4 were treated with 2 μg/mL doxycycline for 48 h to induce mRuby3-RAB5^Q79L^ expression and then observed by time-lapse fluorescence microscopy at 4-s intervals under normal growing conditions (see also Movies S2–S4). Arrowheads indicate ferritin–NCOA4 condensates in the cytosol, and arrows the ferritin–NCOA4 condensates in enlarged endosomes. Scale bars, 5 μm. **(D, E)** 3D-CLEM of cells expressing mGFP-FTH1 treated as in (C). Correlative scanning electron microscopy (SEM) and fluorescent images are shown. An enlarged endosome containing mGFP-FTH1 puncta is magnified. Scale bars, 5 μm (main) and 1 μm (magnified) (D). TEM images of the enlarged endosome indicated by arrows in SEM and fluorescent images are shown. Scale bars, 5 μm (SEM and fluorescent images), 1 μm (TEM), and 100 nm (magnified TEM) (E). **(F)** *NCOA4* KO cells expressing mGFP-FTH1 were examined as in (C). Scale bar, 10 μm.

We also examined the involvement of endosomal microautophagy. To enlarge endosomes so that we could see intraluminal vesicles (ILVs) by fluorescence microscopy, we took advantage of mRuby3-RAB5^Q79L^, a constitutively active mutant of RAB5 (Stenmark et al., 1994; Mejlvang et al., 2018). After doxycycline-induced expression of mRuby3-RAB5^Q79L^, some GFP-FTH1 puncta were seen trapped in enlarged endosomes even under normal growth conditions (Figure 4C, upper panels). Similar results were obtained when mGFP-NCOA4 was used (Figure 4C, middle panels). The magnified images showed that the fluorescence intensity of the GFP-FTH1 puncta inside the enlarged endosomes (Figure 4C, lower panels, arrows) was similar to that in the cytosol (Figure 4C, lower panels, arrowhead), suggesting that ferritin condensates were directly incorporated into endosomes. These puncta in enlarged endosomes moved quickly (Movies S2–S4). Some diffuse GFP signals were detected inside endosomes, likely representing the disruption of ILV membranes (Figure 4C). In addition, 3D-CLEM using scanning electron microscopy (SEM) revealed that the enlarged endosomes containing mGFP-FTH1 puncta were electron-dense and contained electron-dense ILVs (Figure 4D). By TEM, ferritin particles were detected in these ILVs (Figure 4E). In contrast to WT cells, *NCOA4* KO cells did not incorporate mGFP-FTH1 into enlarged endosomes at all (Figure 4F), suggesting that the formation of NCOA4-mediated condensates is required for the incorporation of ferritin into endosomes. Collectively, these observations demonstrate that ferritin–NCOA4 condensates are targeted not only by macroautophagy but also by endosomal microautophagy.

### TAX1BP1 is dispensable for the formation of ferritin–NCOA4 condensates but required for their recognition by macroautophagy and microautophagy

We further examined whether TAX1BP1, an adaptor protein required for ferritin turnover (Goodwin et al., 2017), was involved in ferritin–NCOA4 condensate formation. Fluorescence microscopy showed that mGFP-TAX1BP1 co-localized with mRuby3-FTH1 puncta in WT and *FIP200* KO cells (Figure 5A), suggesting that TAX1BP1 is also a component of ferritin–NCOA4 condensates. These FTH1^+^TAX1BP1^+^ condensates often associated with FTH1^−^TAX1BP1^+^ condensates, which were likely SQSTM1 bodies as we previously observed (Kishi-Itakura et al., 2014). However, knockout of TAX1BP1 (Figure S1A) did not impede condensate formation (Figure 5B). These results denote that TAX1BP1 interacts with component(s) of ferritin-NCOA4 condensates, probably with NCOA4 (Goodwin et al., 2017), but is dispensable for the formation of ferritin condensates. Given that ferritin turnover was blocked in the absence of TAX1BP1, as reported previously (Goodwin et al., 2017) (Figure S1B), we assumed that TAX1BP1 functions after the formation of condensates, more specifically, at the recognition step of ferritin–NCOA4 condensates by macroautophagy and/or microautophagy. In WT cells under iron-depleted conditions, Halo-LC3 frequently co-localized with mGFP-FTH1 puncta (Figures 5C, 5D). By contrast, *TAX1BP1* KO cells did not show co-localization of Halo-LC3 with mGFP-FTH1 puncta (Figures 5C, 5D), suggesting that TAX1BP1 is required for the recognition of ferritin-NCOA4 condensates by macroautophagy. Likewise, mGFP-FTH1 puncta were frequently trapped in endosomes in WT cells, but not in *TAX1BP1* KO cells (Figures 5E, 5F). Taken together, we concluded that TAX1BP1 is dispensable for ferritin–NCOA4 condensate formation, but is required for the recognition of ferritin condensates as a common adaptor for macroautophagy and endosomal microautophagy.

**Figure 5.**
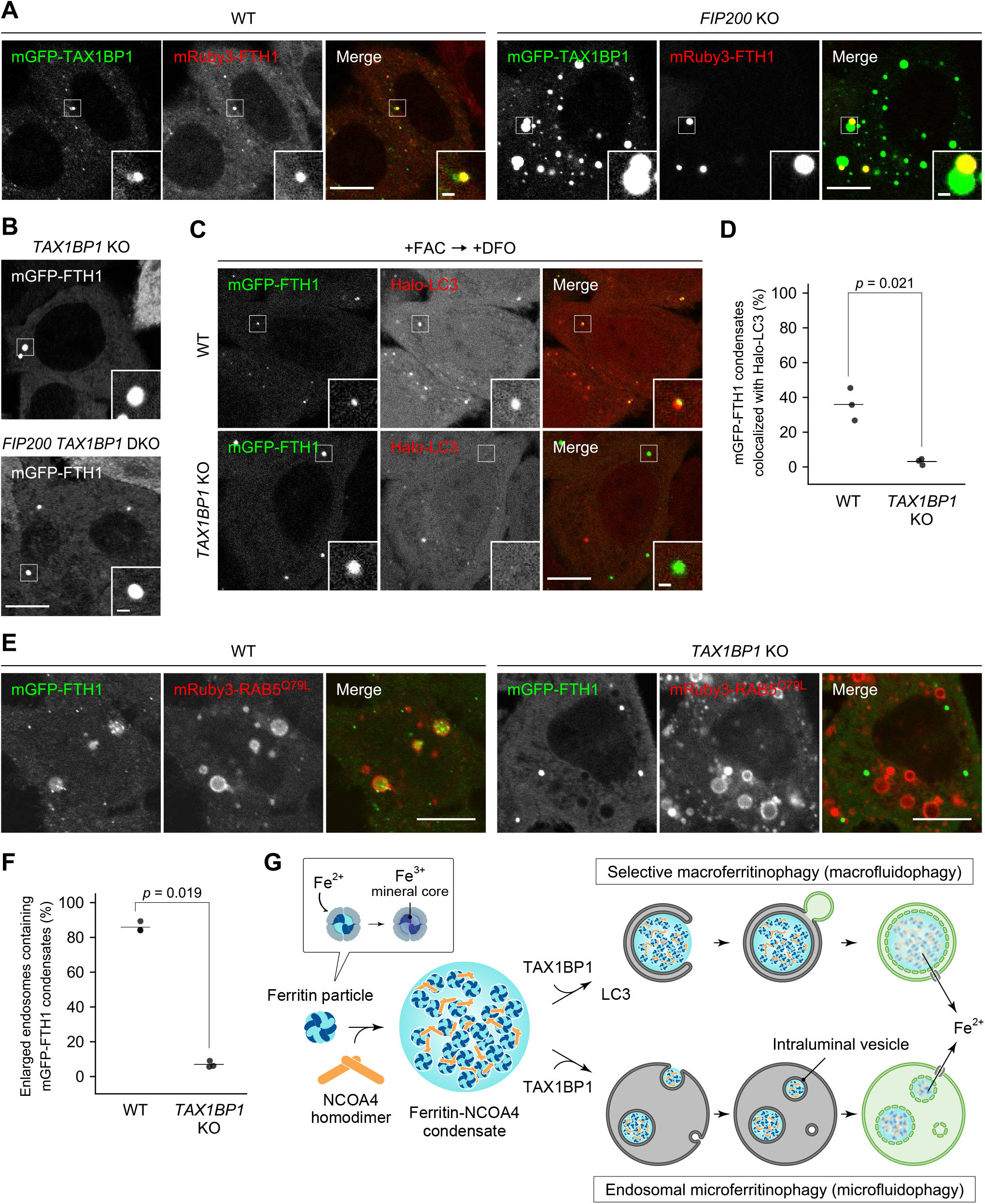
TAX1BP1 is dispensable for ferritin–NCOA4 condensate formation but required for their recognition for macroautophagy and endosomal microautophagy. **(A)** Co-localization of TAX1BP1 with ferritin condensates. WT and *FIP200* KO cells expressing mGFP-TAX1BP1 and mRuby3-FTH1 were grown in DMEM and observed by fluorescence microscopy. Scale bars, 10 μm (main) and 1 μm (inset). **(B)** TAX1BP1 is dispensable for condensate formation. *TAX1BP1* KO and *FIP200 TAX1BP1* DKO cells expressing mGFP-FTH1 were observed as in (A). Scale bar, 10 μm. **(C)** Ferritin–NCOA4 condensates are recognized by autophagosomes in a TAX1BP1-dependent manner. WT and TAX1BP1 KO cells expressing mGFP-FTH1 (green) and Halo-LC3 (red) were treated with 100 μg/mL FAC for 24 h followed by 50 μM DFO for 5.5 h and then observed by fluorescence microscopy. Scale bars, 5 μm (main) and 1 μm (inset). **(D)** The co-localization rate of mGFP-FTH1 puncta with Halo-LC3 in (C) was quantified (*n* = 265–1455). Solid bars indicate the medians, dots indicate the data from three independent experiments. Differences were statistically analyzed by Welch t-test. **(E)** Ferritin–NCOA4 condensates are incorporated into endosomes in a TAX1BP1-dependent manner. mRuby3-RAB5^Q79L^ was expressed by treatment with 2 μg/mL doxycycline for 48 h in WT and *TAX1BP1* KO cells expressing mGFP-FTH1. Scale bars, 10 μm. (F) The rate of the endosomes containing mGFP-FTH1 puncta in (E) was quantified (*n* = 100–198). Solid bars indicate the medians, and dots indicate the data from three independent experiments. Differences were statistically analyzed by Welch t-test. **(G)** A model of NCOA4-dependent formation of ferritin condensates and TAX1BP1-dependent recognition of condensates by macroautophagy and endosomal microautophagy.

### Discussion

In this study, we revealed that ferritin particles undergo phase separation in the cytosol (Figure 5G) and that NCOA4 functions as a driver of ferritin phase separation by providing multivalent interactions (i.e., homodimerization and the direct binding to FTH1) (Figure 3). We also showed that resultant ferritin–NCOA4 condensates were eventually sorted into two different pathways, macroautophagy and endosomal microautophagy (Figure 5G), in a TAX1BP1-dependent manner (Figure 5). Increasing reports have pointed out the relationship between macroautophagy and phase separation of autophagic cargos in various species and contexts, including SQSTM1 bodies in mammalian cells (Sun et al., 2018), Ape1 condensates in the cytoplasm-to-vacuole targeting (Cvt) pathway in yeast (Yamasaki et al., 2020), and PGL granules during embryogenesis of *C. elegans* (Zhang et al., 2018). Recently, it was demonstrated that the autophagosomal sequestration of liquid droplets is promoted by the wetting effect resulting from contact between liquid droplets and membranes (Agudo-Canalejo et al., 2021). This process was termed “fluidophagy,” highlighting the importance of the liquidity of droplets in the deformation of the autophagosomal membranes. During ferritin macroautophagy, we observed that the autophagosomal membranes adhered to the surface of liquid-like ferritin–NCOA4 condensates (Figure 3B), which appears to be comparable to that observed during SQSTM1 fluidophagy (Agudo-Canalejo et al., 2021). Thus, we propose that ferritin macroautophagy (macroferritinophagy) is a type of “macrofluidophagy.” It is possible that ferritin condensates with a liquid-like property promote the elongation of autophagosomal membranes along the surface of ferritin condensates to achieve highly selective degradation.

In addition to ferritin macrofluidophagy, we found that ferritin–NCOA4 condensates were also incorporated into endosomes, suggesting that LLPS promotes cargo sorting into endosomes for lysosomal degradation. We assume that the endosomal microautophagy of ferritin condensates can be regarded as “microfluidophagy,” which may also be promoted by the abovementioned wetting effect. If this is the case, LLPS might be a common mechanism for the two distinct lysosomal ferritin transport pathways. Although selective sorting of microRNAs into exosomes by LLPS of RNA-binding protein has been reported (Liu et al., 2021), little is known about the relationship between LLPS and the endosomal sorting and microautophagy pathways. It is well known that ferritin is secreted and this secretion is enhanced in some diseases, including hemophagocytic lymphohistiocytosis and adult-onset Still’s disease (Rosario et al., 2013). Moreover, autophagy-related proteins such as NCOA4 were found to be secreted via extracellular vesicles (Solvik et al., 2021). Furthermore, a recent paper reported that ferritin secretion via extracellular vesicles depends on NCOA4 (Yanatori et al., 2021). Thus, ferritin–NCOA4 condensates may be directed to both lysosomal degradation and secretion after incorporation into endosomes; this needs to be investigated in further studies.

We also distinguished the roles of NCOA4 and TAX1BP1 in ferritin turnover (Figure 5G). NCOA4 has been thought to be an autophagy receptor for ferritin (Mancias et al., 2014; Dowdle et al., 2014), but we discovered its important additional role in acting as a scaffold of ferritin phase separation by providing multivalent interactions (Figure 3). It is possible that the degradation efficiency of the ferritin–NCOA4 condensates are regulated by the expression levels and ratio of NCOA4, FTH1, and FTL. In fact, the expression levels of FTH1 and FTL respond to iron concentration (Theil, 1987; Munro, 1990), and the FTH1:FTL ratio differs among organs (Arosio et al., 1976). On the other hand, TAX1BP1 is required for the recognition of ferritin condensates rather than condensate formation. TAX1BP1 has a noncanonical LC3-interacting region at residues 141–143 (Newman et al., 2012; Tumbarello et al., 2015) and binds to LC3 family proteins, which play a central role in selective autophagy (Birgisdottir et al., 2013; Johansen and Lamark, 2020). Thus, the association of ferritin–NCOA4 condensates with autophagosomes can be explained by the direct binding of TAX1BP1 to LC3 family proteins. However, the mechanism by which TAX1BP1 links ferritin–NCOA4 condensates to endosomal microautophagy has yet to be elucidated. This pathway may not require LC3 binding given that lysosomal degradation of NCOA4 is partially independent of the LC3 lipidation machinery (Goodwin et al., 2017; Mejlvang et al., 2018). An as yet unidentified factor might be involved in the endosomal microautophagy of ferritin–NCOA4 condensates.

Ferritin clusters have been observed in several types of cells under physiological conditions, including reticulocytes (Sullivan et al., 1976; Heynen and Verwilghen, 1982), Caco-2 cells (Meyron-Holtz et al., 2014), and human kidney proximal tubule brush border cells (Cohen et al.) as well as in some disease conditions such as in erythroblasts in sideroblastic anemia (Ghadially, 1975) and in hepatocytes in hemochromatosis (Iancu, 1992). Further studies are needed to confirm that these structures are indeed ferritin–NCOA4 condensates and are degraded by macroautophagy and/or endosomal microautophagy. We also observed the engulfment of ferritin–NCOA4 condensates by autophagosomes under iron-deficient conditions (Figures 4, 5) and incorporation into endosomes under normal growth conditions (Figures 4, 5), both of which require TAX1BP1 as an adaptor protein. However, the mechanism by which the sorting of ferritin–NCOA4 condensates into the two different pathways (or three if the exosome pathway is included) is regulated remains unclear and needs to be elucidated in the future. Cellular iron metabolism is a network of many reactions and pathways that require various kinds of molecules, including proteins, inorganic iron, and RNAs (Pantopoulos et al., 2012; Bogdan et al., 2016). Biomolecular condensates can function as an organization hub that couples different reactions (Shin and Brangwynne, 2017). Further studies will be required to reveal which molecules exist and what kind of reactions occur in ferritin–NCOA4 condensates so that we can gain an understanding of their exact role in iron metabolism.

## Acknowledgements

We are grateful to Maya Shirakawa for technical assistance with the cell biology experiments; Satoru Takahashi, Yoko Ishida, and Keiko Igarashi for technical assistance with the 3D-CLEM experiments; Shoji Yamaoka for the pMRXIP plasmid; and Teruhito Yasui for the pCG-gag-pol and pCG-VSV-G plasmids. This work was supported by the Exploratory Research for Advanced Technology (ERATO) research funding program of the Japan Science and Technology Agency (JST) (JPMJER1702 to N.M.) and a Grant-in-Aid for Transformative Research Areas (A) from the Japan Society for the Promotion of Science (JSPS) (21H05256 to H.Y.).

## Author Contributions

T.O., H.Y., and N.M. designed the project. T.O. and H.Y. performed the cell biology experiments. Y.S. and C.S. performed the TEM and 3D-CLEM experiments. T.O., H.Y., and N.M. wrote the manuscript. All authors commented on the manuscript.

## Materials and Methods

### Cell lines and culture conditions

HeLa and HEK293T cells (authenticated by RIKEN) were cultured in Dulbecco’s modified Eagle medium (DMEM) (D6546; Sigma-Aldrich) supplemented with 10% fetal bovine serum (FBS) (173012; Sigma-Aldrich) and 2 mM L-glutamine (25030-081; GIBCO) in a 5% CO_2_ incubator at 37°C. For the iron-replete conditions, HeLa cells were treated with 10, 50, or 100 μg/mL ferric ammonium citrate (FAC) (F5879; Sigma-Aldrich). For iron-deficient conditions, cells were treated with FAC for 24 h followed by 50 μM deferoxamine (DFO) (D9533; Sigma-Aldrich).

*FIP200* KO HeLa cells have been described previously (Tsuboyama et al., 2016). *NCOA4* KO, *FIP200 NCOA4* DKO, *TAX1BP1* KO, and *FIP200 TAX1BP1* DKO HeLa cells were generated as follows: DNA fragments encoding the neomycin-resistant cassette flanked by 500-bp sequences homologous to exon 4 and exon 6 of the *NCOA4* gene or homologous to exon 4 of the *TAX1BP1* gene were prepared. WT and *FIP200* KO HeLa cells were cotransfected with the DNA fragment and the PX459 (Addgene #48139)-based plasmid expressing Cas9 and gRNA (GTCTTAGAAGCCGTGAGGTA for the *NCOA4* gene) or gRNA (GTTCTGTTACGTTACCCATA for the *TAX1BP1* gene) using FuGENE HD (E2311; Promega) for 4 h. The cells were cultured for 5 days in DMEM, treated with 1.5 mg/mL G418 (09380-86; Nacalai Tesque) for 1 week, and the clones were selected. HeLa cells inducibly expressing mRuby3-RAB5^Q79L^ were generated as follows: a DNA fragment encoding mRuby3-RAB5^Q79L^ under the Tet-on promoter with the hygromycin-resistant cassette was flanked by 500-bp sequences homologous to the AAVS1 locus. WT, *NCOA4* KO, and *TAX1BP1* KO HeLa cells were cotransfected with the DNA fragment and the PX459 (Addgene #48139)-based plasmid expressing Cas9 and gRNA (GGGGCCACTAGGGACAGGAT). The cells were cultured for 5 days, treated with 50 μg/mL hygromycin (10687010; Thermo Fisher Scientific) for 1 week, and the clones were selected.

### Plasmids

Plasmids for stable expression in HeLa cells were generated as follows: DNA fragments encoding enhanced GFP, monomeric enhanced GFP (mGFP) harboring A206K mutation, mRuby3 (codon-optimized from Addgene #74252), HaloTag7 (Halo) (G1891; Promega), or 3×FLAG were inserted into the retroviral plasmid pMRX-IP (Kitamura et al., 2003; Saitoh et al., 2003) or pMRX-IB (Morita et al., 2018) by the seamless ligation cloning extract (SLiCE) method (Motohashi, 2017). Then, DNA fragments encoding human FTH1 (NP_002023.2), FTL (NP_000137.2), NCOA4 (NP_001138734.1, isoform 3), or TAX1BP1 (NP_001073333.1, isoform 2) were inserted into the pMRX-IP-based or pMRX-IB-based plasmids by the SLiCE method.

For expression in yeast cells, pRS316-based plasmids expressing monomeric ultra-stable GFP (muGFP) (Scott et al., 2018) or muGFP-NCOA4 were generated as follows: a DNA fragment encoding muGFP or muGFP-NCOA4 was inserted downstream of the GPD promoter of pRS316-GPDpro by the SLiCE method. pRS314-based plasmids expressing FTH1 were generated as follows: a DNA fragment encoding FTH1 was inserted downstream of the GPD promoter of pRS314-GPDpro by the SLiCE method.

For the yeast two-hybrid assay, DNA fragments encoding the N-terminal domain (residues 1–182) of NCOA4 (WT, I56E, or L63R) were inserted into the pGADT7 or pGBKT7 vector by the SLiCE method.

### Stable expression in HeLa cells by retrovirus infection

For preparation of the retrovirus solution, HEK293T cells were transfected with the pMRX-IP-based or pMRX-IB-based retroviral plasmid (Kitamura et al., 2003; Saitoh et al., 2003) together with pCG-gag-pol and pCG-VSV-G using Lipofectamine 2000 (11668019; Thermo Fisher Scientific) for 4–6 h. After the cells were cultured for 2–3 days in DMEM, the retrovirus-containing medium was harvested and filtered through a 0.45-μm filter unit (Ultrafree-MC; Millipore) and added to HeLa cells with 8 μg/mL polybrene (H9268; Sigma-Aldrich). After the cells were cultured for 1 day, selection was performed with 1–2 μg/mL puromycin (P8833; Sigma-Aldrich) or 2–3 μg/mL blasticidin (022-18713; Fujifilm Wako Pure Chemical Corporation).

### RNA interference

Stealth RNAi siRNAs (Thermo Fisher Scientific) were used for RNA interference. HeLa cells were transfected with siLuc (CGCGGUCGGUAAAGTTGUUCCAUUU), siFTH1 #1 (CCAGAACUACCACCAGGACUCAGAG), siFTH1 #2 (CAUGUCUUACUACUUUGACCGCGAU), siFTH1 #3 (AGUCACUACUGGAACUGCACAAACU), and/or siFTL (GCAAAGUAAUAGGGCUUCUGCCUAA) using lipofectamine RNAiMAX (13778150; Thermo Fisher Scientific) for 4 h. Then, the cells were cultured for 46 h (siFTH1 and siFTH1 siFTL) or 66 h (siLuc and siFTL) in DMEM.

### Preparation of whole cell lysates

HeLa cells were harvested by centrifugation at 3,000 × *g* for 1 min at 4°C and lysed with 0.2% *n*-dodecyl-β-D-maltoside (DDM) (14239-54; Nacalai Tesque) in 25 mM HEPES-KOH pH 7.2, 150 mM NaCl, 2 mM MgSO_4_, and 1% protease inhibitor cocktail (P8340; Sigma-Aldrich) for 20 min on ice and then treated with 0.1% benzonase (70664; Millipore). The protein concentrations were determined by a microvolume spectrophotometer (NanoDrop One; Thermo Fisher Scientific). Whole cell lysates were mixed with 2×SDS-PAGE sample buffer and boiled at 98°C for 5 min and the protein concentrations were adjusted with 1×SDS-PAGE sample buffer.

### Co-immunoprecipitation

HeLa cells were solubilized with 0.1% DDM in HNE buffer (25 mM HEPES-KOH pH 7.2, 150 mM NaCl, 2 mM EDTA) containing 1% protease inhibitor cocktail (P8340; Sigma-Aldrich) for 20 min on ice and then centrifuged at 17,700 × *g* for 15 min. The supernatants were incubated with anti-DYKDDDDK/FLAG magnetic beads (017-25151; Fujifilm Wako Pure Chemical Corporation) for 3 h at 4°C. The beads were washed three times with HNE buffer containing 0.05% DDM, and bound proteins were eluted with SDS-PAGE sample buffer at 98°C for 5 min.

### Immunoblotting

Immunoblotting was performed using anti-FTH1 (MA5-32244; Thermo Fisher Scientific), anti-FTL (MA5-32755; Thermo Fisher Scientific), anti-NCOA4 (SAB1409837; Sigma-Aldrich), anti-TAX1BP1 (HPA024432; Sigma-Aldrich), anti-FIP200 (17250-1-AP; Proteintech), anti-HSP90 (610419; BD Transduction Laboratories), anti-GFP (A-6455; Thermo Fisher Scientific), and HRP-conjugated anti-DYKDDDDK/FLAG (015-22391; Fujifilm Wako Pure Chemical Corporation) as primary antibodies, and HRP-conjugated anti-rabbit IgG (111-035-144; Jackson ImmunoResearch) and HRP-conjugated anti-mouse IgG (315-035-003; Jackson ImmunoResearch) as secondary antibodies. SuperSignal West Pico Chemiluminescent Substrate (1856135; Thermo Fisher Scientific) and Immobilon Western Chemiluminescent HRP Substrate (P90715; Millipore) were used to visualize the signals, which were detected by an image analyzer (FUSION SOLO.7S.EDGE; Vilber-Lourmat). Contrast and brightness adjustments were performed using the ImageJ (National Institutes of Health) or Photoshop CC 2019/2020 (Adobe) software.

### Yeast transformation

For exogenous expression of muGFP-NCOA4 and FTH1 in yeast cells, BJ2168 *atg11*Δ *atg17*Δ cells were transformed with pRS316-muGFP or pRS316-muGFP-NCOA4 by the high-efficiency yeast transformation method (Gietz and Schiestl, 2007) and grown on SD (-Ura) plates at 30°C. The cells were further transformed with pRS314 or pRS314-FTH1 and grown on SD (-Ura, -Trp) plates. For the yeast two-hybrid assay, AH109 cells were transformed with the pGBKT7-based plasmid and grown on SD (-Trp) plates. The cells were further transformed with the pGADT7-based plasmid and grown on SD (-Trp, -Leu) plates.

### Fluorescence microscopy

Live-imaging fluorescence microscopy was performed using the FV3000 confocal laser microscope (Olympus) equipped with a 60× oil-immersion objective lens (NA 1.4, PLAPON60XOSC2; Olympus) and a stage top CO_2_ incubator (STXG-IX3WX; Tokai Hit) at 37°C with 5% CO_2_. HeLa cells were grown in a glass-bottom dish (3910-035; Iwaki) and fluorescent images were captured using the FluoView software (Olympus). For observation of Halo-LC3, HeLa cells were incubated with 20 nM HaloTag SaraFluor 650T ligand (GCKA308; Promega) for 15 min before observation. The numbers of punctate structures were counted using Fiji software (National Institutes of Health) (Schindelin et al., 2012).

### Fluorescence recovery after photobleaching (FRAP)

FRAP experiments were performed using the FV3000 confocal laser microscope system (Olympus) at 37°C with 5% CO_2_. Photobleaching of mGFP-FTH1 was achieved using a 488-nm laser with a bleaching time of 55.461 ms. Images were captured at 10-s intervals for 20 min (120 time points).

### Transmission electron microscopy (TEM)

HeLa cells were grown on a Celltight C-1 Celldesk LF coverslip (MS-0113K; Sumitomo Bakelite) in DMEM and fixed with 2.5% glutaraldehyde (G018/1; TAAB) in 0.1 M cacodylate buffer pH 7.4 (37237-35; Nacalai Tesque) for 2 h on ice. Postfixation, embedding, and observation under a transmission electron microscope (H-7100; Hitachi) have been described previously (Tamura et al., 2017).

### 3D-CLEM

For observation of the ferritin condensates engulfed by the autophagosomes, HeLa cells expressing mGFP-FTH1 and Halo-LC3 were grown on a glass-bottom dish with 150-μm grids (TCI-3922-035R-1CS; Iwaki, a custom-made product with cover glass attached in the opposite direction) coated with carbon and 0.1% gelatin as described previously (Maeda et al., 2020) and treated with 20 nM HaloTag SaraFluor 650T ligand (GCKA308; Promega) for 15 min before fixation. For observation of the ferritin condensates incorporated into the endosomes or autophagosomes, HeLa cells expressing mGFP-FTH1 were grown as described above and treated with 2 μg/mL doxycycline for 48 h before fixation. The cells were fixed and observed by the FV3000 confocal laser microscope system (Olympus), and then postfixed, embedded in EPON812 as described previously (Maeda et al., 2020). Serial sections (25 nm thick) were cut by an ultramicrotome (UC7; Leica) and observed by a scanning electron microscope (SEM) (JSM7900F; JEOL) and TEM (JEM-1010; JEOL). CLEM images were constructed using Fiji (National Institutes of Health) (Schindelin et al., 2012) and Photoshop CC 2019/2020 (Adobe) software.

## Figure Legends

**Supplementary Figure S1.**
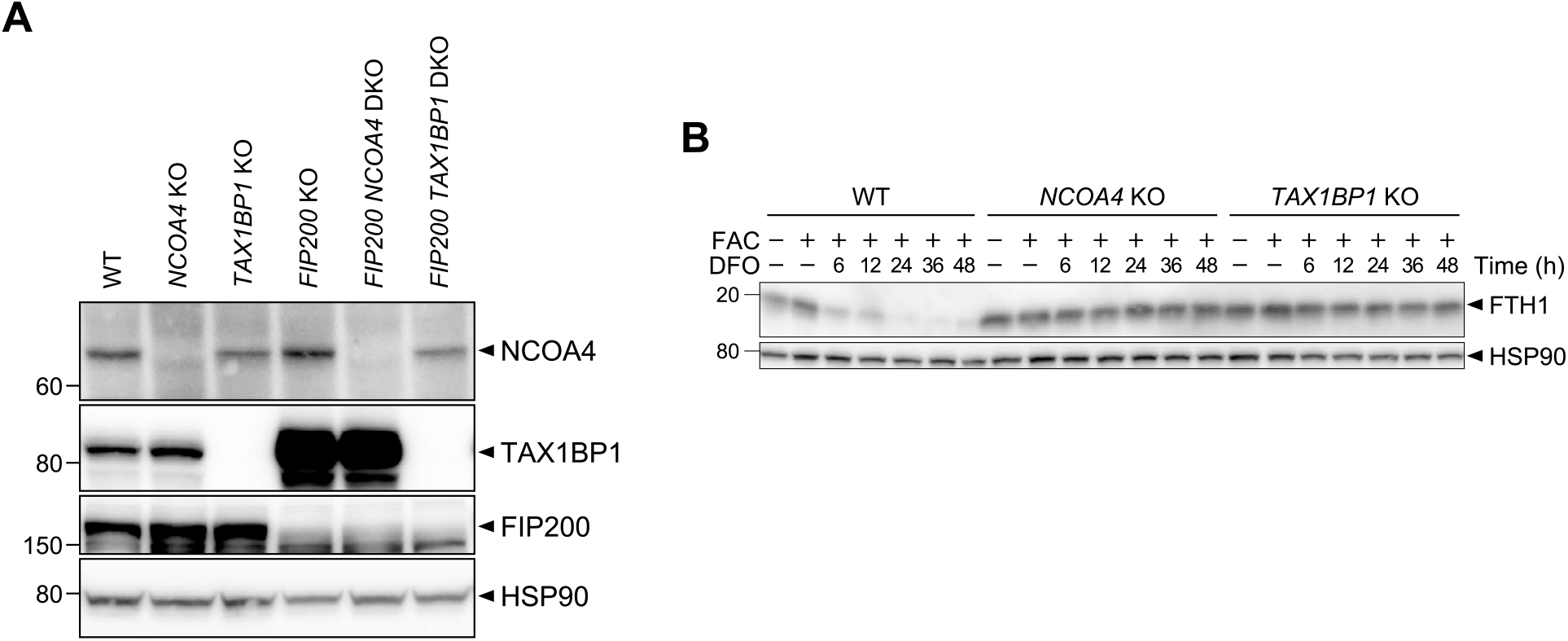
NCOA4 and TAX1BP1 are involved in ferritin degradation. **(A)** WT, *NCOA4* KO, *TAX1BP1* KO, *FIP200* KO, *FIP200 NCOA4* DKO, and *FIP200 TAX1BP1* DKO HeLa cells were grown in DMEM. Whole-cell lysates were analyzed by immunoblotting with antibodies against NCOA4, TAX1BP1, FIP200, and HSP90. **(B)** WT, *NCOA4* KO, and *TAX1BP1* KO cells were grown in DMEM and treated with 10 μg/mL FAC for 24 h followed by 50 μM DFO for the indicated hours. Whole-cell lysates were analyzed by immunoblotting with antibodies against FTH1 and HSP90.

**Supplementary Figure S2.**
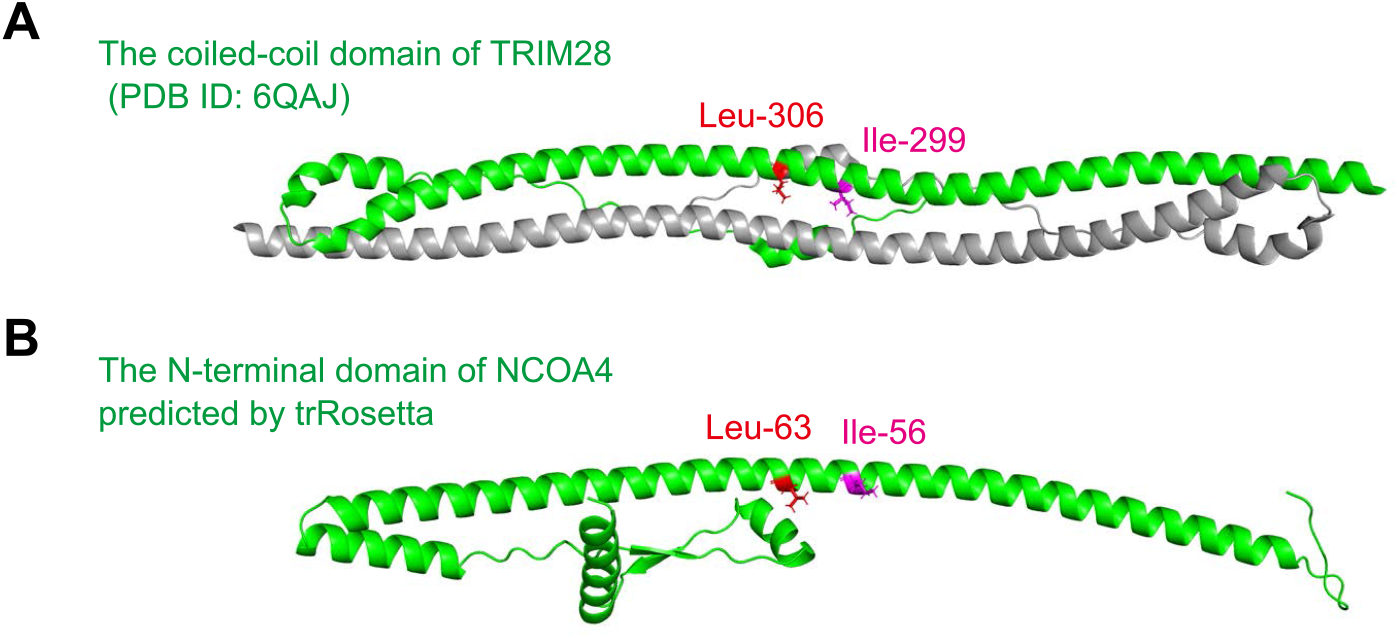
The N-terminal domain of NCOA4 is predicted to form a homodimer. **(A)** An HHpred search showed that the N-terminal domain of NCOA4 is similar to the coiled-coil domain (residues 244–405) of TRIM28, which forms a homodimer. The coiled-coil domains of TRIM28 (green and gray) are shown (PDB ID: 6QAJ). **(B)** The structure of the N-terminal domain (residues 1–182) of NCOA4 is predicted by trRosetta. The putative self-interaction sites Ile-56 (magenta) and Leu-63 (red) are shown.

**Supplementary Figure S3.**
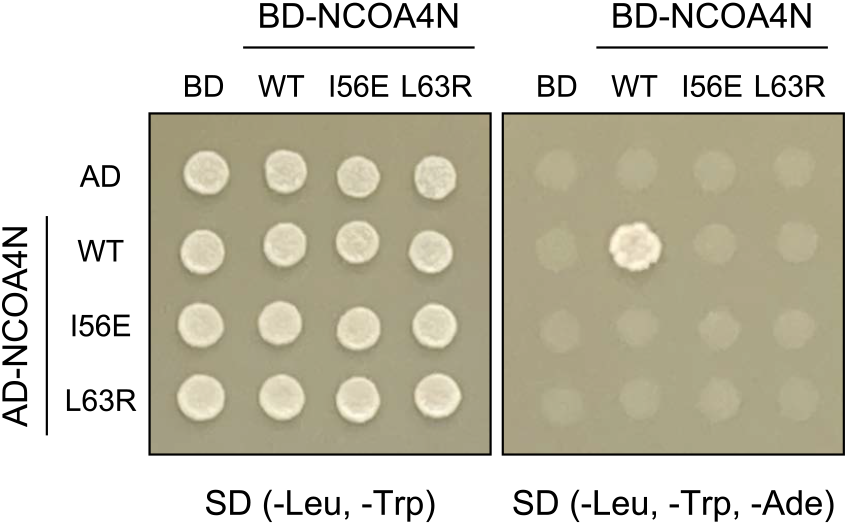
The I56E or L63R mutants of NCOA4 are defective in self-interaction. Yeast AH109 cells were transformed with plasmids expressing the N-terminal domain (residues 1–182) of NCOA4 with or without I56E or L63R mutation fused with a transcription activation domain (AD) or a DNA-binding domain (BD). The cells were grown on SD (-Leu, -Trp) or SD (-Leu, -Trp, -Ade) plates.

**Supplementary Movie S1. A ferritin–NCOA4 condensate is engulfed by an autophagosome**

Fluorescent images of mGFP-FTH1 (green) and Halo-LC3 (red) in WT HeLa cells (used in Figure 4A) were captured at 2-min intervals and are shown at 7 fps.

**Supplementary Movie S2. Ferritin–NCOA4 condensates labeled with GFP-**

**FTH1 are incorporated into enlarged endosomes**

Fluorescent images of GFP-FTH1 in WT HeLa cells expressing mRuby3-RAB5^Q79L^ (used in Figure 4C, upper panels) were captured at 4-s intervals and are shown at 10 fps.

**Supplementary Movie S3. Ferritin–NCOA4 condensates labeled with mGFP-NCOA4 are incorporated into enlarged endosomes**

Fluorescent images of mGFP-NCOA4 in WT HeLa cells expressing mRuby3-RAB5^Q79L^ (used in Figure 4C, middle panels) were captured at 4-s intervals and are shown at 10 fps.

**Supplementary Movie S4. Ferritin–NCOA4 condensates labeled with GFP-FTH1 are observed in the cytosol and an enlarged endosome**

Fluorescent images of GFP-FTH1 in WT HeLa cells expressing mRuby3-RAB5^Q79L^ (used in Figure 4C, lower panels) were captured at 4-s intervals and are shown at 5 fps.

